# Dissecting the Impact of Maternal Androgen Exposure on Offspring Health through Targeting the Androgen Receptor in Developmental Programming

**DOI:** 10.1101/2023.12.12.569558

**Authors:** Haojiang Lu, Hong Jiang, Congru Li, Emilie Derisoud, Allan Zhao, Gustaw Eriksson, Eva Lindgren, Han-Pin Pui, Sanjiv Risal, Yu Pei, Theresa Maxian, Claes Ohlsson, Anna Benrick, Sandra Haider, Elisabet Stener-Victorin, Qiaolin Deng

## Abstract

Women with polycystic ovary syndrome (PCOS) exhibit sustained elevation in circulating androgens during pregnancy, an independent risk factor linked to pregnancy complications and adverse neonatal outcomes. Yet, further investigation is required to understand the precise mechanisms and the impact on cell-type specific placental dysfunction. To explore these dynamics, a PCOS-like mice model was induced with continuous androgen exposure throughout pregnancy, mimicking the human-PCOS. This resulted in impaired placental and embryonic development, leading to mid-gestation lethality. Co-treatment with the androgen receptor blocker, flutamide, prevented this lethality. Comprehensive analysis using whole-genome bisulfite and RNA sequencing revealed the diminished proportion of trophoblast precursors by downregulation of *Cdx2*. The absence of *Gcm1*, *Synb,* and *Prl3b1* further resulted in decreased numbers of syncytiotrophoblasts and sinusoidal trophoblast giant cells, leading to observed compromised placenta labyrinth formation. Importantly, human trophoblast organoids exposed to androgens exhibited analogous alterations, highlighting impaired trophoblast differentiation as a key feature in PCOS-related pregnancy complications. Remarkably, all effects were mediated through the androgen receptor pathways, as demonstrated by comparable offspring phenotypes to controls when treated with flutamide. These findings provide novel insight into the PCOS-related pregnancy complications, and potential cellular targets for future treatment.

## 1. Introduction

Polycystic ovary syndrome (PCOS) manifests in 10-13% of women during reproductive age, giving rise to subfertility marked by irregular menstrual cycles, diminished endometrial receptivity, and endometrial cancer.^[1]^ Additionally, PCOS is associated with comorbidities such as type 2 diabetes, and psychiatric morbidity.^[2]^ Moreover, individuals with PCOS encounter elevated risk of early pregnancy loss, preeclampsia, preterm delivery,^[3]^ and adverse neonatal outcomes.^[4]^ These obstetric challenges confer a predisposition upon offspring to develop reproductive and cardiometabolic disorders in adulthood.^[1, 5, 6]^

The placenta, a transient organ, facilitates the supply of oxygen and nutrients to the developing fetus throughout the entire gestation. Despite variances in physiological and morphological features, both human and mouse placentas exhibit a hemochorial structure, wherein the fetal trophoblast establishes direct contact with maternal blood circulation.^[7]^ The trophoblast lineage is principally composed of cytotrophoblast cells, giving rise to syncytiotrophoblasts in both species, and invasive extravillous trophoblasts in humans along with murine trophoblast giant cells. Syncytiotrophoblasts play a pivotal role in placental steroidogenesis and hormonal production, while invasive trophoblast cells are crucial for the successful establishment of pregnancy.^[8]^ Previous studies have demonstrated that dysfunction in any of these cell types can lead to adverse outcomes such as pre-eclampsia, uterine growth restriction, or even miscarriage.^[7, 9]^ Similarly, restoration of essential placental gene expression, resulting in the replenishment of critical placental cell types, has been shown to prevent embryo lethality.^[10, 11]^

Women with PCOS exhibit sustained elevation in circulating androgens and anti-Müllerian hormone (AMH) levels throughout pregnancy,^[12, 13]^ which has been shown as independent risk factors for pregnancy complications and adverse neonatal outcomes. Notably, women with PCOS manifest an unfavorable maternal-fetal environment characterized by aberrant placenta morphology, diminished invasion sites, and impaired steroidogenesis activities,^[14–16]^ thereby impacting fetal development. Nevertheless, the specific mediation of these adverse effects and the repercussions on distinct placental trophoblast cell types in the context of maternal hyperandrogenism remains further investigation.

We and others have demonstrated that maternal hyperandrogenism exerts deleterious effects on fetal development,^[17, 18]^ consequently predisposing their offspring to subsequent reproductive, metabolic, and psychiatric disorders.^[19–21]^ Specifically, daughters born to women with PCOS exhibit a fivefold increase in the likelihood of being diagnosed with PCOS^[6]^, and are prone to develop psychiatric disorders later in life.^[22]^ Additionally, newborns display an elongated anogenital distance (AGD) a strong marker of *in utero* androgen excess.^[6, 23]^ Conversely, sons born to women with PCOS face an elevated risk of developing obesity, hyperlipidemia, anxiety, and depression.^[5, 24, 25]^ Previous studies show that androgen receptor activation results in altered implantation and uterine mitochondrial dysfunction in androgenized pregnant rats, as well as diminish decidualization and angiogenesis potentially mediating the transmitted effects to the offspring.^[17]^ However, whether and how maternal androgen exposure affect the placenta, early embryo and primordial germ cell development, and whether co-treatment targeting androgen pathways has the potential to prevent placenta dysfunction, results in normal fetal and germ cell development as well as healthy offspring remains to be investigated.

We here present evidence that an *in utero* hyperandrogenic environment in mice profoundly compromises both placental, embryo and germ cell development, with a preventing effect through the administration of the androgen receptor blocker, flutamide. Employing whole genome bisulfite sequencing (WGBS) and RNA sequencing, we identified molecular alterations in primordial germ cells (PGCs) and placentas at critical embryonic developmental time points, specifically embryonic days (E)10.5 and E13.5. We revealed significant impairment in the differentiation of trophoblast cell lineages within the placenta, culminating in miscarriage in female mice exposed to androgens during pregnancy. Additionally, disruptions in placental fatty acid metabolism were identified suggesting a potential predisposition of the offspring to metabolic abnormalities later in life. These adverse effects were determined to be mediated through androgen receptor pathways as the female and male offspring exposed to androgens *in utero* exhibited normal phenotypes upon treatment with flutamide. Moreover, human trophoblast organoids exposed to androgens demonstrated a reduction in trophoblast differentiation and invasion capacity, an effect also ameliorated by flutamide. These findings contribute novel insights into complications associated with PCOS during pregnancy, thereby informing the development of future treatment modalities.

## 2. Results

### 2.1. Flutamide mitigates reproductive dysfunction induced by androgen exposure

A PCOS-like mouse model was established by implantation of dihydrotestosterone (DHT) pellets into 4-week-old peripubertal F0 female mice, hereafter referred to as PCOS-mice, in comparison to control mice implanted with inert pellets. ^[26]^ To investigate the potential causal involvement of the androgen receptor pathway in the development of PCOS-like traits and to assess the preventive capacity of downstream effects, one group of mice received simultaneous implantation of a DHT-pellet and a continuous slow-releasing flutamide pellet, an androgen receptor antagonist, hereafter referred to as flutamide mice (**Fig. 1a**). Phenotypic assessments were conducted in all F0 mice after 10 weeks of exposure.

**Fig. 1.**
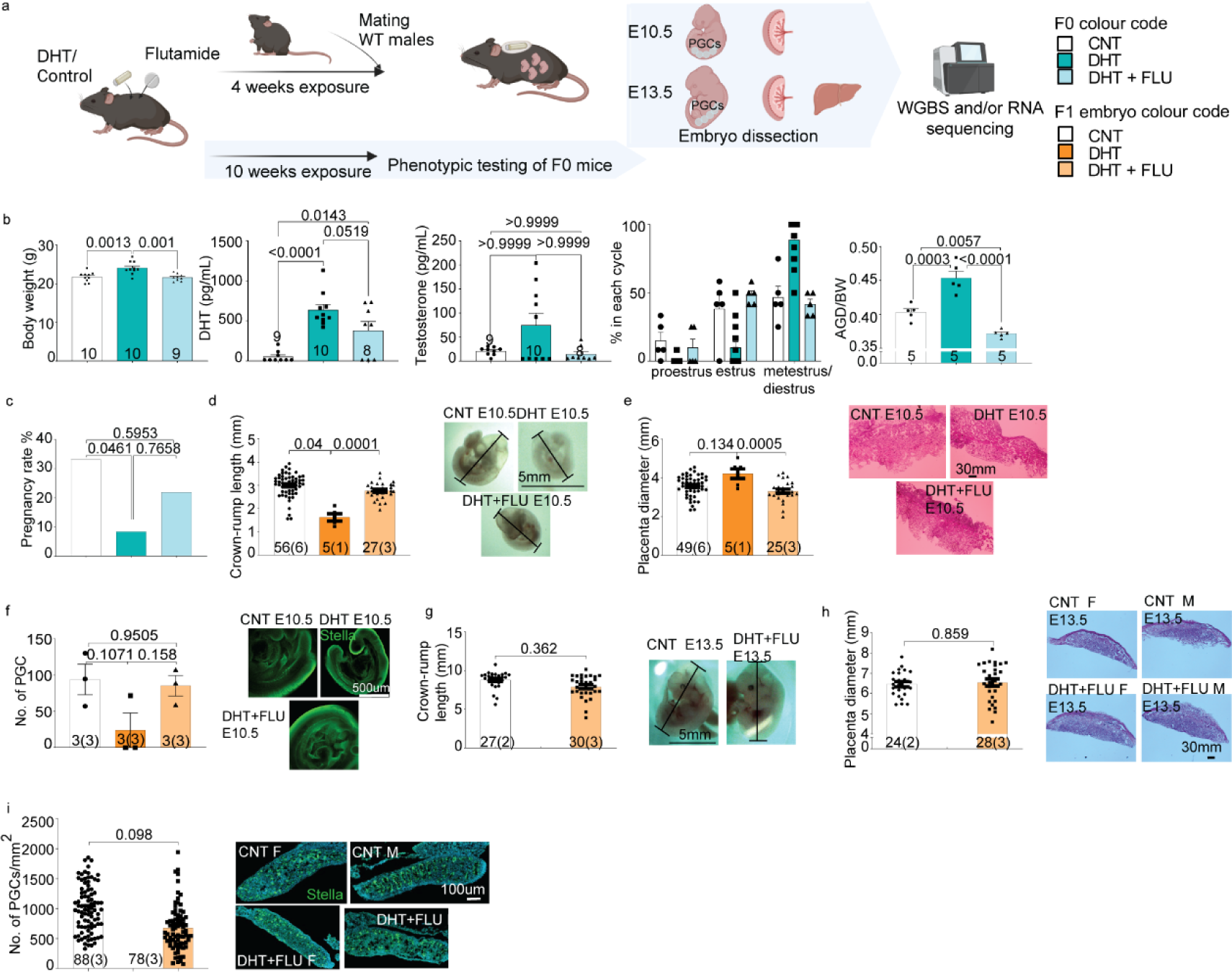
Peripubertal androgen-induced PCOS-like mouse model negatively affects pregnancy outcomes that are prevented by co-treatment with flutamide. **a,** Schematic overview of the experimental design and breeding scheme of the embryonic part. Control = vehicle; DHT = implanted with slow-releasing DHT pellet; DHT + FLU = implanted with DHT pellet and a flutamide pellet. **b,** Peripubertal PCOS-like (F0) mice weigh more (Control = 10, DHT = 10, DHT + FLU = 9), have increased circulating DHT levels measured by LC-MS, disrupted estrus cyclicity (P<0.05 for M/D phase CNT vs DHT and DHT vs FLU comparison; CN T= 5, DHT = 15, DHT + FLU = 5), longer anogenital distance (AGD), and developed typical polycystic ovarian morphology according to Hemotoxilin and Eosin (H&E) stains. **c,** Pregnancy rate and number of embryos per F0 dam; Control = 40, DHT = 40, DHT + FLU =28. **d,** Crown-rump length and representative images of E10.5 embryos from each group. **e,** Placenta diameter and representative bright-field images of H&E staining of E10.5 placentas. **f,** Number of PGCs counted in fixed E10.5 embryos; Control = 3, DHT = 3, DHT + FLU = 3. And representative confocal microscope images of E10.5 whole mount embryo staining of PGCs with Stella. **g,** Crown-rump length, and representative images of E13.5 embryos from each group. **h,** Placenta diameter and representative bright-field images of H&E staining of E13.5 embryo placentas. **i,** Number of primordial germ cells (PGCs) counted in fixed E13.5 embryo gonads; Control = 3/gender, DHT + FLU = 3/gender and representative confocal microscope images of E13.5 gonads with PGCs stained with Stella. All data are presented as mean ± SEM. Numbers of mice are stated in the text of each graph or in the bar of each group with the number of F0 mothers given in the bracket. F0: One-way ANOVA with Turkey post hoc analysis if passed the Shapiro Wilk normality test; otherwise Kruskal-wallis test is used. F0 body weight development: Repeated measures ANOVA. F1 embryos: use of ANCOVA to control for litter size whenever possible otherwise One-way ANOVA with Turkey post hoc analysis was used.

Following 10-weeks of DHT exposure, the PCOS-mice exhibited an increase in body weight compared to controls, while the flutamide-mice were protected against weight gain (one-way ANOVA: *F*_2, 26_ = 10.42; *P* =0.0005) (**Fig. 1b**). Circulating DHT level was higher in both PCOS-mice and flutamide-mice (one-way ANOVA, *F*_2, 24_ = 16.82; *P* =0.0242) (**Fig. 1b**), with no discernable difference in circulating testosterone level between the groups (Kruskal Willis test, *H* = 2.833; *P =* 0.2426) (**Fig 1b**). Reproductive function was evaluated through a 12-day continuous vaginal smear, revealing that the PCOS-mice were consistently in metestrus/diestrus phase indicating complete anovulation. In contrast, control- and flutamide-mice exhibited normal cyclicity (**Fig. 1b**). Elongated AGD, a robust indicator of androgen exposure, was evident in PCOS-mice compared to controls, and this effect was prevented by flutamide treatment (one-way ANOVA: *F*_2,12_ = 38.81; *P* = <0.0001) (**Fig. 1b**). These findings affirm the successful establishment of PCOS-like mouse model through peripubertal DHT pellet implantation and underscore the androgen receptor pathways as mediators of the observed phenotypic traits.

### 2.2. Flutamide mitigates abnormal placental and embryonic development induced by maternal androgen excess

To comprehensively assess the impact of maternal androgen excess on placental function and embryonic development, we used the peripubertal DHT-induced PCOS mouse model with or without flutamide pellets to interrogate the effect of androgen receptors activation. Following 4 weeks of exposure, these mice were mated with wildtype males, necessitating superovulation due to anovulatory feature of the PCOS-mice. A total of 40 PCOS-mice, 40 control female mice (F0), and 28 flutamide-treated PCOS-mice were used for this purpose. Embryos were collected and dissected at E10.5 and E13.5, two critical stages to examine the early- and mid-embryonic development and dynamics of embryo DNA methylation (**Fig. 1a**).

The PCOS-mice (F0) exhibited a diminished pregnancy rate compared to the controls, an effect that was completely prevented by flutamide (Kruskal-Wallis: *H* = 5.882; *P* = 0.041) (**Fig. 1c**). Additionally, the quantification of deceased/absorbed embryos per dam at the time of dissection revealed a higher count in the PCOS-mice, a phenomenon that was also normalized by flutamide co-treatment at both E10.5 and E13.5 (E10.5: Kruskal-wallis test, *H* = 7.017; *P =* 0.0159; E13.5: student t test, *P >* 0.9999) (**Extended Data Fig. 1b**).

In addition to the diminished pregnancy rate observed in the PCOS-mice, embryos were smaller, as indicated by a shorter crown-rump length at E10.5. Notably, this effect that was absent in the flutamide-treated group (ANCOVA: *P*=0.04) (**Fig. 1d**). PCOS-mice exhibited a trend towards larger placenta compared to control and flutamide groups (ANCOVA: *P_CNT vs DHT_* = 0.134, *P_DHT vs DHT + FLU_* = 0.0005) (**Fig. 1e**). Morphological examination of the PCOS-mice placenta at E10.5 revealed aberrant organization of the labyrinth layer characterized by a smaller labyrinth area and disorganized trophoblast giant cells. In contrast, the flutamide-mice showed a comparable placental morphology to that of the control-mice (**Fig. 1e**). Additionally, the embryos from PCOS-mice exhibited a smaller number of migrating PGCs at E10.5, albeit insignificant, it was prevented by flutamide treatment (ANCOVA: *P* = 0.1071) (**Fig. 1f**). Whole-mount staining of the embryo at E10.5 also revealed a delay in both embryo and PGC development, evident from the migratory PGC location (**Fig. 1f**). All embryos from PCOS-mice failed to develop beyond E13.5. Consequently, only embryos from control and flutamide-treated dams were available for analyses at E13.5. Notably, E13.5 embryos from the flutamide-mice were similar to those of controls, with no difference in crown-rump length, placental size, the number of migrating PGCs, or the placenta morphology in both sexes (**Fig. 1g-i**). Collectively, these findings underscore the pivotal role of androgen receptor pathway activation in mediating maternal effects in PCOS.

### 2.3. Progeny derived from PCOS-dams treated with flutamide exhibit normal development

Next, we aimed to elucidate how a hyperandrogenic *in utero* environment transmits reproductive, metabolic, and anxiety-like behavioral phenotypes to offspring at 4 months and 6 months of age (**Fig. 2a, 3a**). Due to the miscarriage observed in the PCOS-mice at mid-gestation (**Fig 1c**), we decided to use a 5 mm DHT implant.^[26]^ These mice develop a PCOS-like phenotype akin to the 10 mm DHT-implant, characterized by increased body weight, elongated AGD, and impaired glucose tolerance, all of which were effectively prevented by flutamide treatment (**Extended Data Fig. 1a**). Despite the reduced DHT dose used, and after 4 different attempts involving a total of 55 control mice, 115 PCOS-mice and 44 flutamide-treated PCOS-mice, the yield of PCOS-offspring was notably limited (3 females from 2 dams and 9 males from 3 dams) (**Supplementary Table. 1**). Given the resultant lack of statistical robustness, PCOS-offspring are excluded from subsequent analyses to avoid potential misinterpretations.

**Fig. 2.**
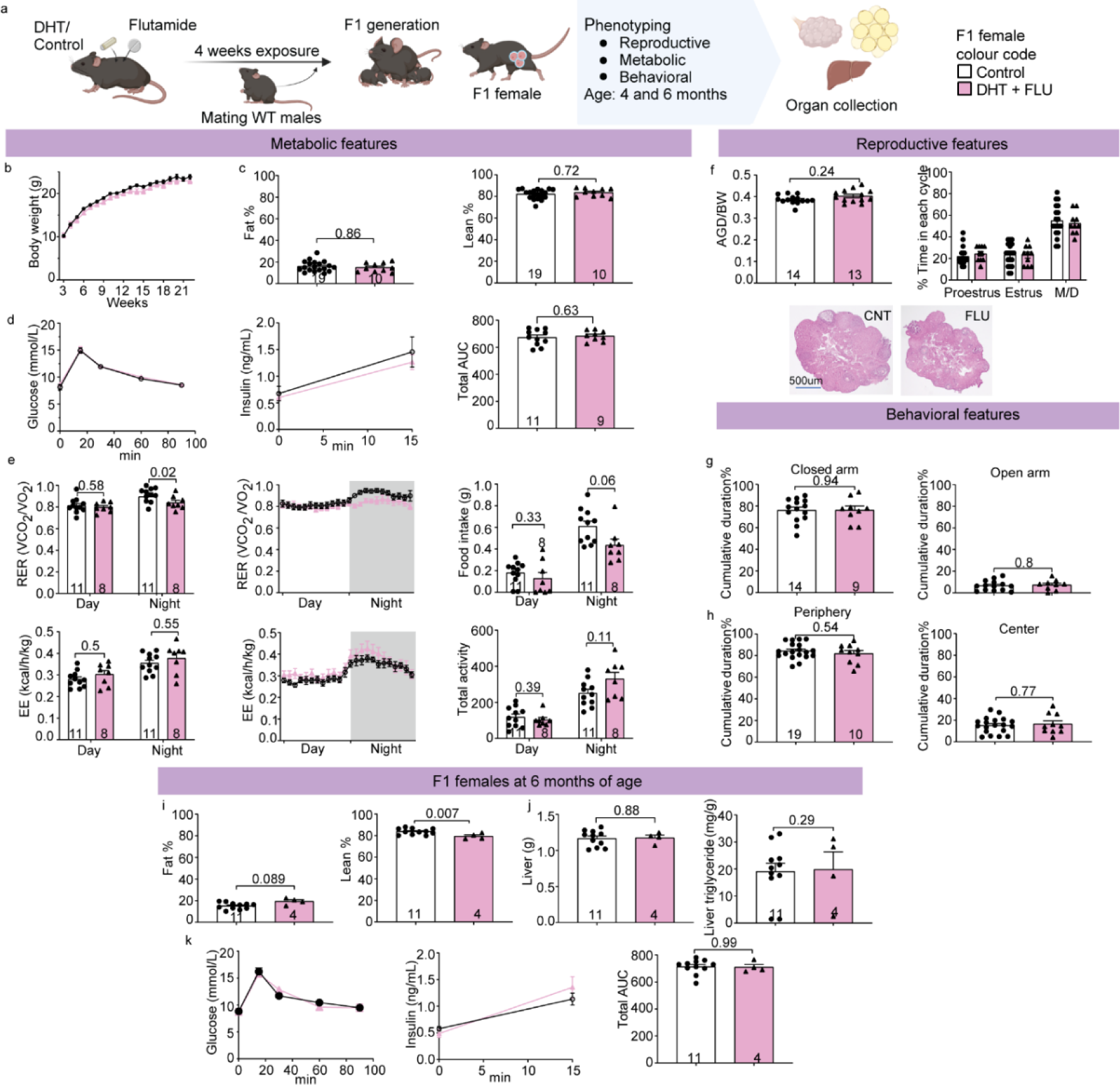
Flutamide treatment prevented the development of any disease phenotypes in F1 female offspring. **a,** Schematic overview of the offspring experimental design and breeding scheme. **b,** Body weight development in F1 female offspring; Control offspring = 16, DHT + FLU = 10. **c,** Percent of fat and lean mass (EchoMRI) normalized to body weight of 4 months age F1 female offspring**. d,** Glucose and insulin curve during oral glucose tolerance test (OGTT) and total area under the curve (AUC) calculation at 4 months of age in F1 female offspring. **e,** Metabolic cage results for F1 females at 4 months of age, including Respiratory Exchange Ratio (RER), Energy Expenditure (EE), Food intake, and total locomotor activity. Grey area indicates nighttime (18 pm to 6 am). **f,** Reproductive phenotype measures for F1 females at 4 months of age, including anogenital distance (AGD) normalized by body weight, estrus cyclicity (Control offspring = 19, DHT + FLU = 10), and representative bright-field images of ovarian morphology. **g,** Time spent in the closed and open arms during elevated plus maze test for 4 months age F1 female offspring. **h,** Time spent in the periphery and center region during open field test for 4 months age F1 female offspring. **i,** Percent of fat and lean mass (EchoMRI) normalized to body weight of 6 months age F1 female offspring. **j,** Dissected liver weight and liver triglyceride level measurement of F1 female after termination of experiment (>6 months of age). **k,** Glucose and insulin curve during OGTT and total AUC calculation of 6 months of age F1 female offspring. All data are presented as mean ± SEM. Numbers of mice are stated in the bars of each group. Body weight development statistics are done by repeated measurement ANOVA. F1 statistics: ANCOVA to control for litter size.

In 4-months-old female and male offspring derived from control and flutamide-mice, no discernible difference was observed in terms of body weight development (**Fig. 2b**, **Fig. 3b**), body composition (**Fig. 2c, 3c**), or glucose and insulin profiles according to the oral glucose tolerance test (OGTT) (**Fig. 2d, 3d**). However, flutamide-female offspring exhibited a slight reduction in respiratory exchange ratio at night (ANCOVA, *P* = 0.02) (**Fig. 2e**), while other variables, including energy expenditure, food intake and total activity, remained comparable between the groups in both female and males (**Fig. 2e**, **Fig. 3e**). Female offspring from flutamide-treated dams displayed no difference in AGD while a decreased AGD was observed in male offspring (**Fig. 2f**, **Fig. 3f**). In addition, estrus cyclicity, ovarian morphology (**Fig. 2f**), ovary weight, and the number of oocytes (**Extended Data Fig. 1c**) in female offspring were comparable to those of female controls. Similarly, testis morphology (**Fig. 3f**), testis weight, and the number of motile sperms after swim-up assay (**Extended Data Fig. 1d**) in male offspring were also restored. Behavioral phenotypic testing revealed no difference in elevated plus maze test (**Fig. 2g**, **Fig. 3g**) and open field test (**Fig. 2h**, **Fig. 3h**) between flutamide-treated and control-offspring, both in females and males. Reassessment at 6 months of age indicated that, although flutamide-treated female offspring exhibited slightly lower lean mass, no difference was observed in fat mass (**Fig. 2i**). Similarly, there was no difference in body composition (**Fig. 3i**), liver weight, liver triglyceride level (**Fig. 2j**, **Fig. 3j**), glucose, and insulin profiles (**Fig. 2k**, **Fig. 3k**) between female and male offspring. Taken together, the blockade of maternal androgen-receptor pathways during pregnancy successfully mitigated the adverse effect of DHT exposure during fetal development, resulting in a normal reproductive and metabolic phenotype in adulthood.

**Fig. 3.**
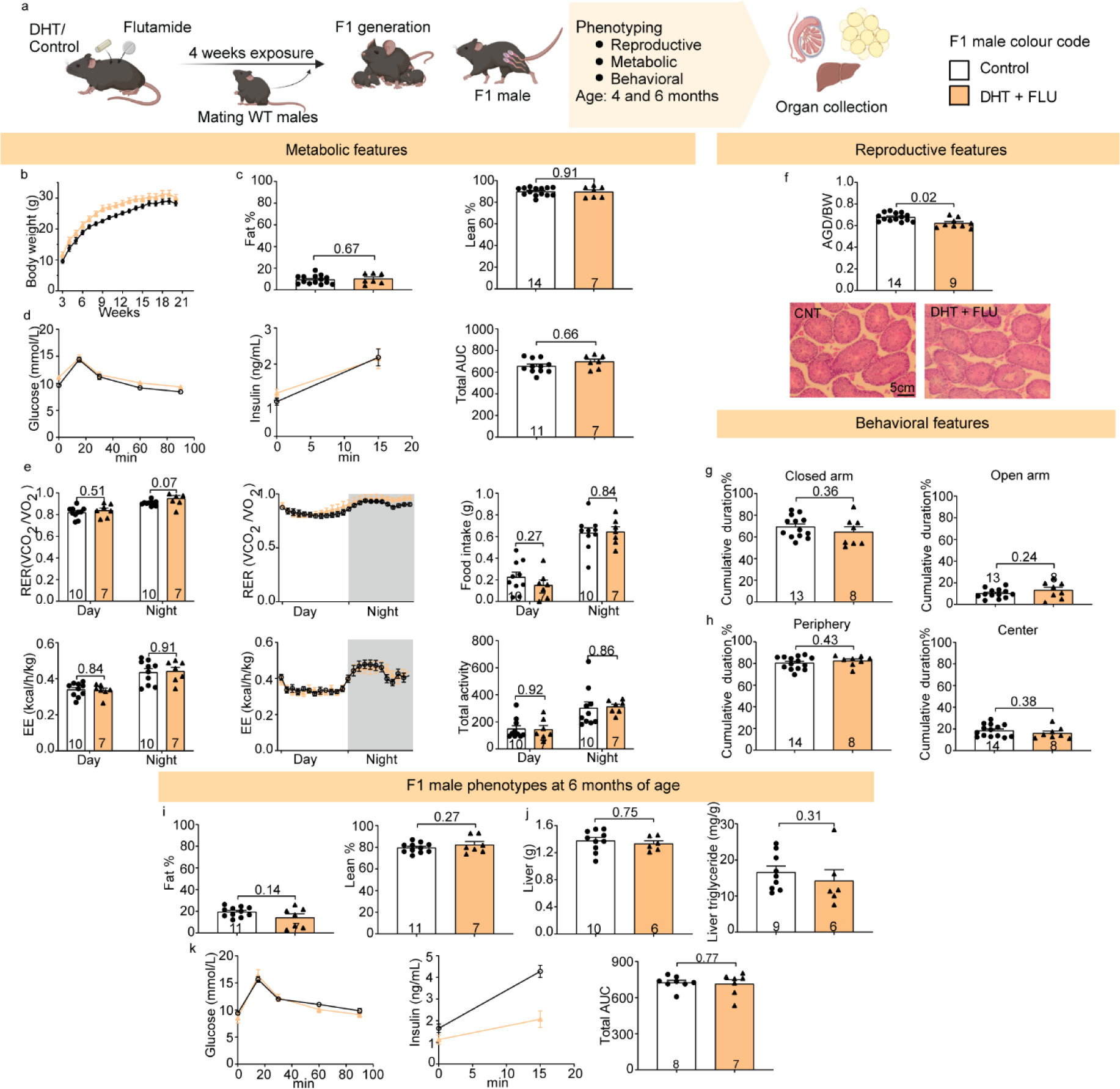
Flutamide treatment prevented the development of any disease phenotypes in F1 male offspring. **a,** Schematic overview of the offspring experimental design and breeding scheme. **b,** Body weight development in F1 male offspring; Control offspring = 13, DHT + FLU = 8. **c,** Percent of fat and lean mass (EchoMRI) normalized to body weight of 4 months age F1 male offspring. **d,** Glucose and insulin curve during oral glucose tolerance test (OGTT) and total area under the curve (AUC) calculation at 4 months of age in F1 male offspring. **e,** Metabolic cage results for F1 males at 4 months of age, including: Respiratory Exchange Ratio (RER), Energy Expenditure (EE), Food intake and total locomotor activity. Grey area indicates nighttime (18pm to 6am). **f,** Reproductive phenotype measures for F1 males at 4 months of age, including: Anogenital distance (AGD) normalized by body weight and testis morphology. **g,** Time spent in the closed and open arms during elevated plus maze test for 4 months F1 male offspring. **h,** Time spent in the periphery and center region during open field test for 4 months age F1 male offspring. **i,** Percent of fat and lean mass (EchoMRI) normalized to body weight of 6 months age F1 male offspring**. j,** Dissected liver weight and liver triglyceride level measurement of F1 males after termination of experiment (>6 months of age). **k,** Glucose and insulin curve during OGTT and total AUC calculation at 6 months of age in F1 male offspring. All data are presented as mean ± SEM. Numbers of mice are stated in the bars of each group. Body weight development: Repeated measures ANOVA. F1: ANCOVA whenever possible to control for litter size.

### 2.4. Androgen exposure disrupts trophoblast proliferation and differentiation in the placenta

Subsequently, we investigated the molecular profile of placentas derived from PCOS-mice compared to controls and flutamide-treated mice (**Extended Data Fig 2a, Supplementary Tables 2 and 3**). At E10.5, a total of 783 genes showed decreased expression, while 435 genes exhibited increased expression (**Fig. 4a, Extended Data Fig 2a, b, Supplementary Table 4**). Notably, placentas from flutamide-mice displayed a transcriptional profile indistinguishable from that of control mice at both at E10.5 and E13.5, indicating that the inhibition of the androgen receptor safeguarded against the transcriptional perturbation observed in PCOS-mice (**Fig. 4a, Extended Data Fig 2a**).

**Fig 4.**
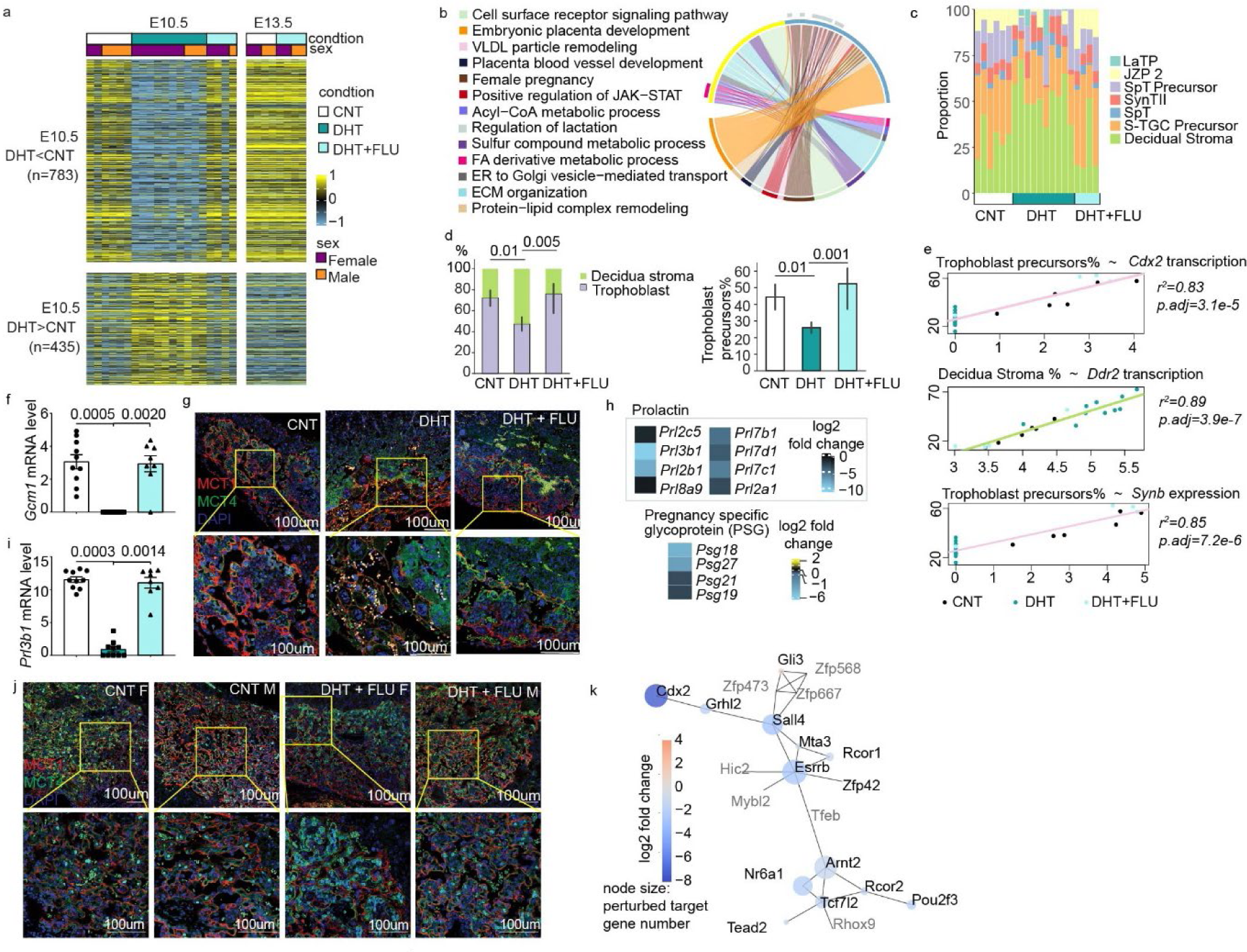
Flutamide prevented placenta development abnormalities from PCOS-mice at E10.5. a, heatmap displaying the expression pattern of 1218 DEGs (differentially expressed genes) in PCOS-mice compared to control-mice at E10.5. The color bar indicates the scaled log2 normalized counts. b, selected cellular process gene ontology pathways enriched by DEGs in PCOS-mice compared to control-mice at E10.5. The pathways shown are enriched using clusterProfiler package (v4.0.5). In the upper half of the sphere, the yellow region are up-regulated genes, and the blue are the down-regulated genes. The lower half shows the enriched pathways. The arcs link the genes to their related pathways c, Cell type proportions deconvoluted by CIBERSORTx. d, Bar plots representing mean proportion of trophoblast cells and trophoblast precursors. The error bars indicate 95% confidence intervals. P values by ANCOVA to control for sex. e, linear regression of selected DEGs and trophoblast or trophoblast precursors proportion. P values are adjusted using Holm method. f, *Gcm1* mRNA levels from RNA sequencing at E10.5. g, Confocal microscopy images of E10.5 mouse placenta stained for syncytiotrophoblast layer I and II (SynTI and SynTII) respectively with MCT1, MCT4 and DAPI; upper row images were at 10X magnification and lower row at 20X magnification. h, Gene expression level from RNA sequencing at E10.5. i, *Prl3b1* expression from RNA sequencing at E10.5. j, Representative confocal microscopy images of E13.5 mouse placenta stained for SynTI and SynTII respectively with MCT1, MCT4 and DAPI; upper row images were at 10X magnification and lower row at 20X magnification. k, Co-expression network of differentially expressed transcription factors in placentae from DHT-mice. The edges indicate the co-expression (median confidence >=0.4). The node size indicates the number of its target genes which were DEGs in PCOS-mice (min: 0, max: 965). The grey gene names were those without a record in TFlink database, thus, they got zero sizes.

Enriched Gene Ontology Biological Process revealed that maternal androgen exposure disrupts pathways associated with female pregnancy and placental development (**Fig. 4b, Extended Data Fig 2c**). At E10.5, the mouse placenta undergoes a significant transformation, marking the shift from the primitive choriovitelline configuration to the chorioallantoic configuration. This transformation involves the expansion of the embryonic labyrinth and the progressive differentiation of trophoblast cell lineages.^[27]^ To understand the impact of maternal androgen excess on placental development and cell type composition, we deconvoluted our bulk RNA-seq expression data using the cell type specific annotation from single-nuclei data at E10.5.^[28]^ The placentas from PCOS-mice exhibited an increased proportion of decidua stromal cells but decreased proportion of trophoblast cells with a nearly 50% less trophoblast precursor cells compared to the control and flutamide-treatment (**Fig. 4c-d, Supplementary Table 5**).

The reduction of trophoblast precursors may be due to the drastic decrease in the expression of stem cell maintenance genes, such as *Cdx2,* which positively correlated with the percentage of trophoblast precursors that was significantly reduced in PCOS-placenta (**Fig.4d-e**). The gene expression of *Ddr2*, encoding the discoidin domain receptor tyrosine kinase 2 and responsible for sensing the extracellular matrix (ECM) collagens,^[28]^ was positively correlated with decidua stroma which was increased in the PCOS-mice placenta (**Fig. 4e**). Indeed, the ECM including collagens was elevated in PCOS-mice placenta (**Fig. 4b**), which could impede the proper proliferation and invasion of trophoblasts.^[29]^ Consistent with these findings, the differentiation of trophoblast precursor cells into syncytiotrophoblast was disturbed with the lowest expression of *Synb* observed in PCOS placentas since *Synb* showed a positive correlation with trophoblast precursor cells (**Fig. 4e**). Indeed, the PCOS-mice placenta retained more LaTP (labyrinth trophoblast precursor) cells instead of differentiating into SynTI (syncytiotrophoblast layer I) and SynTII (syncytiotrophoblast layer II) branches (**Extended Data Fig. 2d**). The *Aqp3* (Aquaporin 3) plays a role in transporting glycerol, free fatty acids, and triglyceride and subsequently influence fetal growth in mice,^[29]^ and the expression of *Aqp3* correlated highly with the LaTP cells arrest (**Extended Data Fig. 2e**). These findings suggests that excess maternal androgens cause dysregulation of proliferation and differentiation of trophoblast precursors cells, and consequently affecting trophoblast fusion.^[30]^

To validate the cell type-specific defects, immunofluorescent staining of trophoblast precursor cells and syncytiotrophoblast cells was performed on E10.5 and E13.5 placentas. In mouse placentas, syncytiotrophoblast precursors undergo terminal differentiation before fusing to form two layers of syncytium: Syn I and Syn II. This process is regulated by glial cells missing transcription factor 1 (*Gcm1*), syncytin A (*SynA*)^[31]^ and syncytin B (*SynB*).^[32, 33]^ Maternal androgen exposure resulted in depletion of *Gcm1* (**Fig. 4f**) and *SynB* (**Extended Data Fig. 2f**), leading to substantial reduction in the formation of Syn I and Syn II layers in E10.5 placenta (**Fig. 4g**). Consequently, a decrease in the expression of gene families of prolactin and pregnancy specific glycoproteins (PSGs) in PCOS-mice placenta was observed, as these are secreted by trophoblast (**Fig. 4h**).^[34]^ In addition, the labyrinth zone patterning in PCOS-mice placentas was altered, with increased trophoblast giant cells (TGCs) population compared to control placentas (**Fig. 4g**).

Although morphologically there was increased presence of trophoblast giant cells in the PCOS-mice placentas at E10.5, the proportion of sinusoidal-TGCs was lower (**Fig. 4c**), along with reduction in the sinusoidal-TGC markers *Prl3b1* and *Ctsq* (**Fig. 4i, Extended Data Fig. 2f**). In line with placenta transcriptome analysis, flutamide treatment effectively prevented the aforementioned placental abnormalities (**Fig. 4f-g, 4i**). PCOS-mice showed embryonic lethality before E13.5, possibly attributed to the deformation of the labyrinth zone and dysregulation of critical genes in placental development.^[35, 36]^ Further analyses of the formation of placental labyrinth zone, especially Syn I and Syn II layers at E13.5, revealed that placentas from flutamide-treated PCOS-mice closely resembled those of controls (**Fig. 4j**).

Interestingly, when comparing DEGs at E10.5 to their expected regulation during normal pregnancy progression at E13.5, we found that most of the DEGs at E10.5 were regulated in the opposite direction at E13.5 (**Extended Data Fig. 2g**). Specifically, genes involved in the differentiation from LaTP to SynII cells in the labyrinth and the invasive sinusoidal-TGC were down-regulated in PCOS-mice, contrary to the expected to be up-regulation at E13.5 (**Extended Data Fig. 2h**), indicating that maternal androgen exposure impedes the labyrinth differentiation.

The co-expression network analysis of differentially expressed transcription factors in PCOS-mice revealed the central role of oestrogen-related receptor beta (*Esrrb)* in transcriptional regulation. *Esrrb* expression was downregulated in PCOS-mice placentas, together with several other transcription factors (**Fig. 4k**). Consistent with previous findings where *Esrrb* function as a key transcription factor involved in the maintenance of trophoblast stem cells,^[37, 38]^ the reduction of *Esrrb* in PCOS-placentas may, in part, explain the phenotypes mentioned above. These also align with the previously observed overexpansion of trophoblast giant cells and impaired labyrinth zone formation in *Esrrb* knockout mice.^[39]^ Maternal hyperandrogenism has been associated with offspring psychiatric illness,^[40]^ association analysis of the the Mamalian Phenotype Ontology^[41]^ with the down regulated DEGs in E10.5 DHT placenta confirmed the implication of *Esrrb* in abnormal brain development (**Supplementary Table 6,7**). This underscores the pivotal role of *Esrrb* as one of the main targets influenced by a hyperandrogenic *in utero* environment.

### 2.5. Fatty acid metabolism regulated by PPAR signalling pathway is disrupted in the PCOS-placentas

Given the increased adiposity observed in PCOS, we investigated whether such alteration could affect the placenta development. Metabolic processes, specifically fatty acid metabolism and the associated Acyl-CoA metabolism, were found to be elevated in placentas from PCOS-mice (**Fig. 4b**, **Fig. 5a**). The upregulation of *Acsl1*, encoding an enzyme responsible for synthesizing fatty acyl-CoAs from long-chain-fatty-acid,^[42]^ and *Nudt7,* encoding enzyme involved in the hydrolysis of fatty acyl-CoAs,^[27]^ was observed in PCOS-mice (**Fig. 5b**). Nonetheless, the protein−lipid complex remodelling pathway was downregulated in PCOS-mice placentas (**Fig. 5a**). ^[42]^ Specifically, *Hmgcs2*, encoding HMG-CoA synthase, involved in a metabolic pathway providing lipid-derived energy,^[43]^ exhibited a four-fold increase in expression in PCOS-mice compared to controls (**Fig. 5b**).

**Fig. 5.**
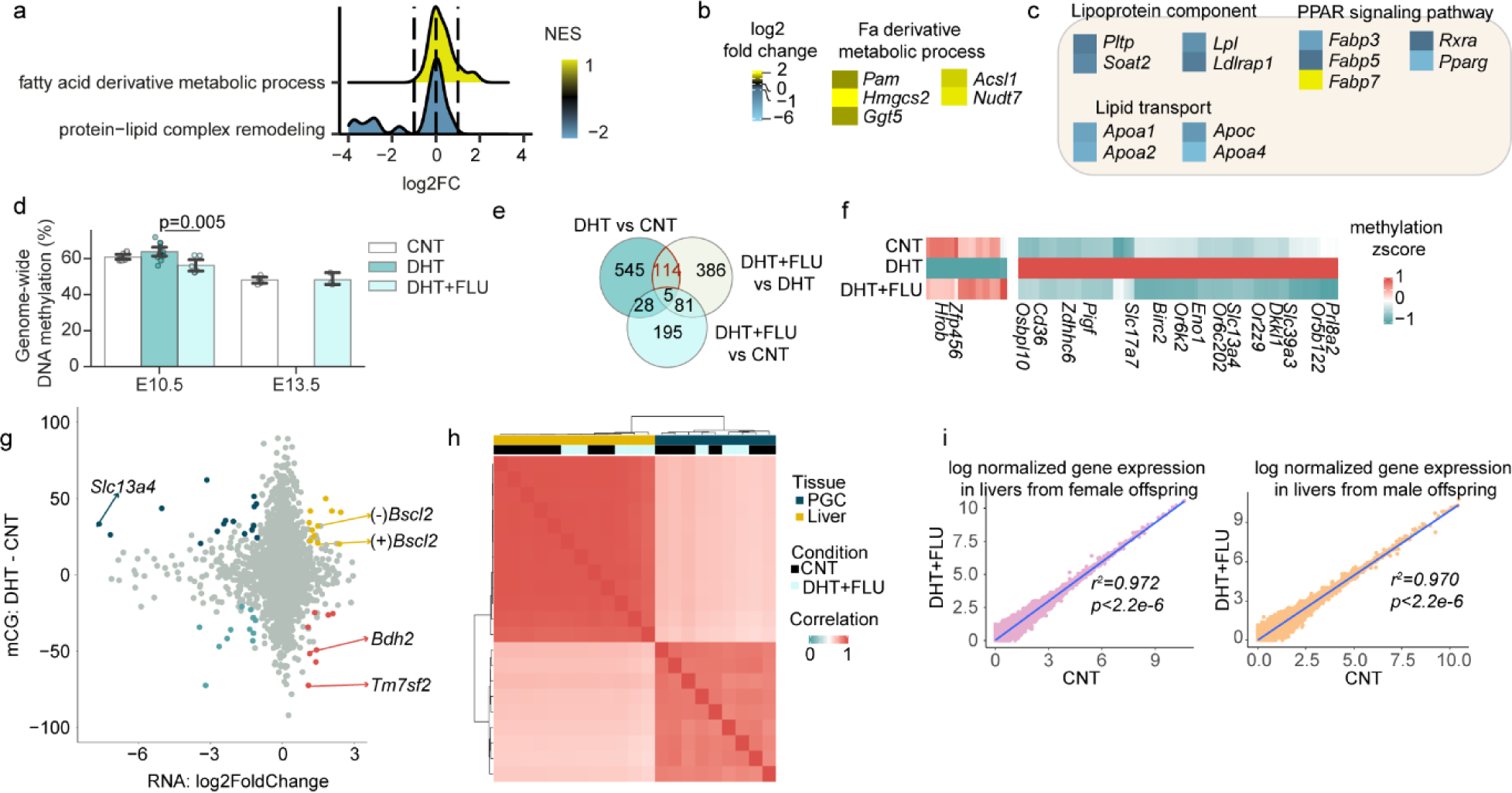
Flutamide co-treatment protects against DNA methylation disturbances in placenta induced by androgen exposure while maintaining the DNA transcription in PGC, embryonic and offspring liver tissues. **a,** The gene set enrichment with the distribution of log2 fold change and normalized enrichment scores. **b,** Transcription analysis of E10.5 CNT vs PCOS placenta DEGs involved in GO term ‘fatty acid derivative metabolic process’. **c,** Downregulated gene expression of DEGs involved in lipid transport, lipid storage (lipoprotein component), and PPAR (Peroxisome proliferator-activated receptor) signaling pathway. The gene set of each metabolic process to intersect with the DEGs is from KEGG database and the enrichment is by a hypergeometric test with q-value < 0.05. Data are presented as log2 fold change, and all genes displayed were DEGs by DESeq2. **d,** Bar plots representing 5mC (5-methylcytosine) level. The error bars indicate 95% confidence intervals. P values by ANCOVA to control for sex. **e,** Veen diagram showing the intersection of the genes annotated with the differentially methylated promoters in each comparison. **f,** The methylation z-score ((methylation level of each sample – mean value of methylation levels)/SD of methylation levels) of the genes whose promoter methylation was dysregulated by DHT treatment and reversed by DHT and FLU cotreatment. **g,** In placenta at E10.5, the differentially expressed genes by DHT treatment and the 5mC sites methylation difference on their promoters. The genes with differentially methylated sites (FDR<5%, difference>20) were colored. **h,** Correlation heatmap of the gene expression in E13.5 PGC and embryonic liver. **i**. Scatter plot and correlation of the gene expression in offspring livers from control and flutamide-mice.

In parallel, a gene set enrichment analysis of pathways from the KEGG database confirmed a decreased enrichment of lipid storage, transport via lipoprotein, and upstream PPAR signalling pathway in placenta from PCOS-mice compared with controls (**Fig. 5c**). These findings suggest a high turnover of acyl-CoA and subsequent lipogenesis from fatty acid oxidation in response to androgen exposure in the placenta. Furthermore, the decrease in lipoprotein components suggested that lipids generated as a response to androgen exposure were not stored as lipoprotein but used for transport.

### 2.6. Flutamide treatment prevented the methylation induced by maternal androgen exposure

The methylation of promoter regions has been associated with alterations in gene transcription,^[30]^ and is therefore investigated in the placenta. The establishment and maintenance of global DNA methylation were only marginally increased by androgen exposure at E10.5 (**Fig. 5d**.). To specifically examine the 5mC (5-methylcytosine) sites in gene promoters, we conducted genome-wide DNA methylation profiling to identify androgen exposure-associated 5mC methylation alterations. At E10.5, androgen exposure led to a slight increase in global 5mC levels in the placenta (ANCOVA, P = 0.25), while flutamide placenta maintained the 5mC level at both E10.5 and E13.5 like the control placenta (ANCOVA, P_CNT vs DHT + FLU_ = 0.11) (**Fig. 5d**).

By quantifying the number of genes with promoters exhibiting distinct 5mC sites between groups, we identified 692 genes associated with PCOS-mice compared with controls, 586 genes associated with PCOS-mice compared with flutamide-mice, and only 309 in flutamide-mice compared with controls (FDR <5%, methylation difference > 20%) (**Fig. 5e**). The differentially methylated 5mC sites were mainly located within CpG shores, known to be susceptible to methylation changes in tissue specification and certain disease progress (**Extended Data Fig 2i**).^[44, 45]^ Moreover, the androgen exposed placentas also affected methylated sites in the promoter region, which were associated with a higher number of DEGs compared with flutamide-mice The flutamide-mice placentas reversed the 5mC methylation changes on promoters associated with 114 genes (**Fig. 5e**). Among them, *Slc13a4* was hypermethylated in the promoter of PCOS-mice but remained unaffected in flutamide-mice (**Fig. 5f**). This hypermethylation might contribute to the 7-fold downregulation of *Slc13a4* transcription in PCOS-mice placenta (**Fig. 5g**). *Slc13a4* encodes the most abundant sulfate transporter located at the syncytiotrophoblast, and the *Slc13a4* loss-of-function in mouse placentas led to fetal death.^[46]^ These findings suggest that the mid-gestation lethality in PCOS-mice resulted from both transcriptional and epigenetic aberrations.

Despite flutamide preventing adverse effects on both placenta and embryo development, concerns arise regarding its long-term effects on offspring health, especially the potential hepatotoxicity.^[47]^ Germ cells are essential in transmitting parental information to the offspring in mice. At E10.5, we collected migrating PGCs for RNA sequencing and DNA methylation profiling. Despite a limited number of samples, the transcriptome of PGCs from embryos of flutamide-treatment was comparable to that of controls (**Extended Data Fig 2j**). Based on the DNA methylation of all 5mC sites, PGCs from embryos of flutamide-treated mice clustered with those of controls but were separated from the PCOS-mice (**Extended Data Fig 2k**). At E13.5, the gene expression profile of germ cells and embryonic livers remained similar in flutamide-mice compared to controls (correlation >0.94, **Fig. 5h**). In the embryonic liver, there were no differentially expressed genes between the control and flutamide-treated groups. While gender is the primary driver for expression differences in livers at 6-month-old (**Extended Data Fig. 2l**), the gene expression patterns were preserved in the livers of both female and male offspring of flutamide-mice (**Fig. 5i**).

### 2.7. Androgen exposure affects villous cytotrophoblast differentiation in human trophoblast organoids

To further explore the translatability of these findings to humans, we established human trophoblast organoids (TOs) cultured from trophoblast stem cells and comprising cytotrophoblast layer and syncytiotrophoblast layers. Extravillous cytotrophoblast, formed upon differentiation, will migrate through the matrigel, leaving a visible trace for observation.^[48]^ Control (CNT group), 1nM DHT (DHT group), and 1nM DHT +2uM flutamide co-treatment (DHT + FLU group) were included in the study. No discernible difference in the appearance, morphology, or size of TOs were observed between CNT, DHT or DHT + FLU groups (Kruskal-wallis test, *H* = 8.884; *P* = 0.0118), and all groups produced human chorionic gonadotrophin (hCG) to a level detectable by commercial pregnancy tests (**Fig. 6a**). Immunofluorescent staining of cytotrophoblast and syncytiotrophoblast markers using EPCAM and ENDOU, respectively revealed that both the CNT and DHT + FLU groups exhibited a typical TO structure. However, TOs treated with DHT had less syncytiotrophoblast and an increased number of cytotrophoblast cells in the centre of the TOs (**Fig. 6b**). Upon induction of differentiation, both CNT and DHT + FLU TOs showed extensive migration of extravillous trophoblasts into the matrigel, while less migration was observed in TOs treated with DHT (**Fig. 6c**). These results validated our findings in the mouse placentas and suggest that androgen exposure profoundly influenced the differentiation of human trophoblast precursors and the invasion of trophoblast cells.

**Fig. 6.**
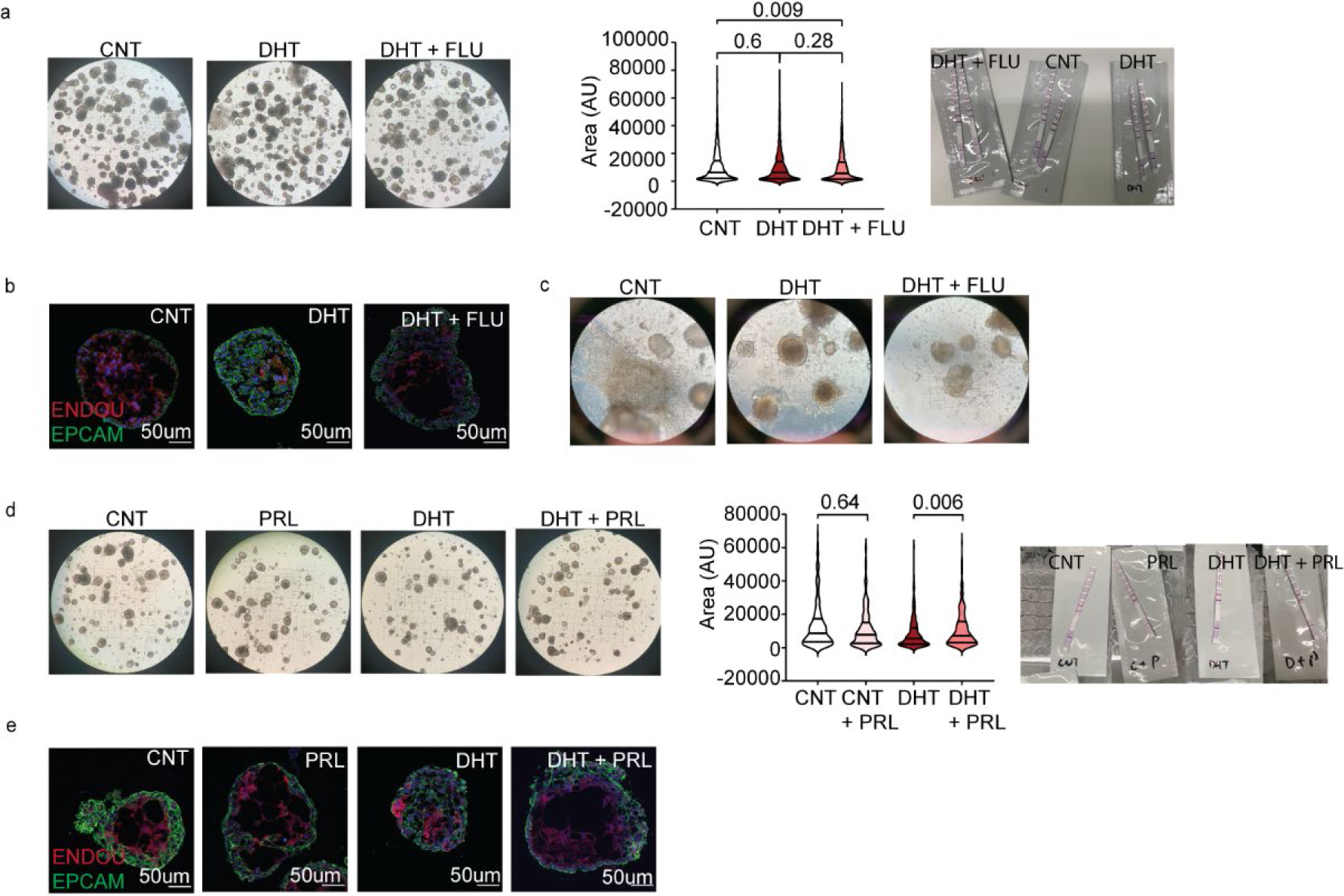
Effect of DHT on mouse placenta and human placenta trophoblast organoids are prevented by flutamide treatment. **a,** Representative images of human trophoblast organoids (TOs) grown in matrigel upon treatment of DHT and/or flutamide, images were taken at 4X magnification. And size measurement of TOs and detection of human chorionic gonadotrophin (hCG) production with commercial pregnancy tests. **b,** Representative confocal microscopy images of organoids stained for cytotrophoblast and syncytiotrophoblast with EPCAM, ENDOU and DAPI, images were taken at 20X magnification. **c,** Representative images of human placenta organoids grown in matrigel after 10 days of differentiation treatment, images were taken at 10X magnification. **d,** Representative images of human placenta organoids grown in matrigel upon treatment of DHT and/or prolactin, images were taken at 4X magnification. And size measurement of TOs and detection of hCG production with commercial pregnancy tests. **e,** Representative confocal microscopy images of organoids stained for cytotrophoblast and syncytiotrophoblast with EPCAM, ENDOU and DAPI, images were taken at 20X magnification. TO size measurement: Kruskal wallis test due to data sets are non-parametric.

Considering the severe downregulation of the prolactin family in E10.5 PCOS-mice placenta (**Fig. 4h**), we next explored whether prolactin could prevent the adverse effects induced by androgen exposure, given its previously demonstrated role in enhancing differentiation and invasion of human trophoblast cells.^[49]^ The prolactin experiment consisted of the following groups: control (CNT group), 100ng/ml prolactin (PRL group), 1nM DHT (DHT group), 1nM DHT + 100ng/ml prolactin (DHT + PRL group). Prolactin did not alter the overall morphology and hCG production in TOs, with or without DHT. However, TO size was increased in the DHT + PRL group compared to the DHT-only group (Kruskal-wallis test, *H* = 22.22; *P* < 0.0001) (**Fig. 6d**). Immunofluorescence staining showed that prolactin treatment improved syncytiotrophoblast differentiation in the DHT + PRL group compared to the DHT group (**Fig. 6e**) without affecting the normal trophoblast differentiation in PRL group. Collectively, these findings suggest that PRL had the potential to promote cytotrophoblast differentiation in placentas under PCOS condition.

## 3. Discussion

Women with PCOS are at increased risk of pregnancy complications, and their progeny are predisposed to an altered growth trajectory and an elevated likelihood of acquiring a PCOS diagnosis, along with associated comorbidities during adulthood.^[50]^ The precise mechanism mediating the transmission of the disease via the maternal-fetal environment remains unexplored. Using the peripubertal DHT-exposure model, ^[50]^ we could analyze a hyperandrogenic maternal environment throughout the entire gestation mimicking human PCOS pregnancy. Specifically, we sought to determine whether continuous maternal androgen exposure-induced alterations could be mitigated through targeted treatment of the androgen receptor, utilizing flutamide. Indeed, the induction of a classic PCOS-like reproductive and metabolic phenotype in F0 dams was successfully counteracted by blocking the androgen receptor pathway, in line with previous research.^[17, 18]^ Moreover, *in utero* hyperandrogenism resulted in a global reduction of the transcriptional activity in the placentas leading to diminished placental labyrinth formation as well as impaired proliferation and differentiation of trophoblast precursors. Consequently, syncytialization was severely compromised, potentially contributing to defective nutrient transport and embryonic lethality at mid-gestation. These deleterious effects were completely prevented by inhibition of the androgen receptor pathway with flutamide, resulting in fetal development and offspring health comparable to controls. Analogous findings were observed in the human trophoblast organoid experiment, indicating conserved effects of hyperandrogenism in disrupting maternal-fetal interaction.

The maintenance of trophoblast stem cell population is critical for supporting its differentiation into different trophoblast cell lineages supporting placenta function.^[51]^ The markedly reduced trophoblast precursors at E10.5 in PCOS-mouse placentas are consistent with previous finding with reduced proliferation of cytotrophoblasts in women with PCOS.^[52]^ This observed reduction might be attributed to a significant downregulation of genes that contributes to maintaining trophoblast stem cells, including *Esrrb*, *Cdx2* and *Elf5*. Notably, the deletion of either *Elf5* or *Esrrb* causes embryonic lethal during early or mid-gestation in mice.^[53, 54]^ Therefore, these genes could be the driver of the embryonic lethality and placental dysfunction observed in PCOS-mice.

One vital step of pregnancy establishment in species with the hemochorial placenta involves proliferation and differentiation of trophoblast stem cells together with the invasion of trophoblasts into the maternal decidua for the remodeling of spiral arteries, a process orchestrated by trophoblast giant cells (TGCs) in mice and extravillous trophoblasts in humans.^[8, 55]^ The formation of syncytiotrophoblast and sinusoidal-TGCs, two important cellular components of the labyrinth zone was severely compromised under maternal hyperandrogenic environment. Sinusoidal-TGCs separate the maternal sinusoids and syncytiotrophoblast layers and are in direct contact with maternal blood circulation facilitating maternal-fetal connection and nutrient transfer.^[56]^ Indeed, genetic ablation of sinusoidal-TGCs is associated with intrauterine growth restriction and embryo death at late-gestation.^[57]^ The great decrease in sinusoidal-TGC marker, *Ctsq*, expression in placentas coincides with the miscarriage in PCOS-mice.

Furthermore, the finding that androgen exposure impedes the invasion of extravillous trophoblasts was confirmed in human trophoblast organoids, suggesting a potential link to miscarriage in the PCOS-mice and women with PCOS. Trophoblast giant cells, known for their endocrine functions, secrete prolactin in mouse placenta.^[58]^ Prolactin is an important hormone involved in angiogenesis and the regulation of immune responses.^[59]^ The substantial downregulation of nine members of the prolactin family, with expression decreases ranging from1-10-fold in androgen-exposed mouse placentas, may have detrimental effects on embryo survival, leading to observed mid-gestation embryo lethality. While the role of prolactin in the human placenta is less explored, it has been suggested to improve trophoblast migration and invasion and to decrease placental inflammation.^[49, 60]^ Consistent with previous findings, our observations revealed improved cytotrophoblast differentiation into syncytiotrophoblast in human trophoblast organoids under a hyperandrogenic environment when supplemented with prolactin, implying a protective role of prolactin in maternal-fetal interaction.

Another critical aspect of pregnancy maintenance involves nutrient and gas transport via the placenta, which is mediated via syncytiotrophoblasts in both mice and humans.^[61]^ The differentiation of syncytiotrophoblasts from trophoblast stem cells requires *Gcm1* and syncytin in mice and humans. *Gcm1* promotes trophoblast stem cells to syncytiotrophoblasts and extravillous trophoblasts differentiation,^[62]^ while syncytin induces cell-cell fusion for the formation of syncytiotrophoblast layers.^[32, 33]^ The 6-to-7-fold decreased expression in both genes contributed to disruption of trophoblast stem cells to syncytiotrophoblasts differentiation thus malformation of syncytiotrophoblast layers. To date, no studies have investigated the impact of PCOS on syncytiotrophoblasts in the human placenta, although a reduced number of syncytiotrophoblasts has been associated with *in utero* growth restriction and pre-eclampsia, both of which are observed in PCOS pregnancies.^[63, 64]^ We showed that DHT exposure impaired syncytiotrophoblast differentiation in human trophoblast organoids, confirming the role of impaired syncytiotrophoblast lineage in PCOS pregnancies.

Fatty acid uptake in trophoblast cells is facilitated by Acsl-mediated conversion into acyl-CoA, which is used for subsequent metabolic processes like β-oxidation.^[65]^ Although maternal androgen exposure increases placental fatty acid metabolism and acyl-CoA uptake, there was concurrent increase in *Nudt7* and *Hmgcs2* expression. This may lead to diminished levels of CoA derivatives in the placenta, thereby inducing mitochondrial stress, inflammation, and pre-eclampsia.^[66]^ Additionally, diminished FABP (fatty acid binding protein), PPARɤ and its functional unit RXRɑ expressions were all observed in the PCOS placenta. FABPs bind fatty acids in the cytoplasm and transport them to their final destinations, including to the nuclear receptors, PPAR and RXR.^[67]^ Considering that the activation of PPARɤ-RXR heterodimer positively regulates fatty acid uptake in human trophoblasts,^[68]^ we hypothesized that maternal hyperandrogenism results in restricted fatty acids transport at the maternal-fetal interface. This limitation could lead to a constrained energy supply for fetal development, potentially resulting in embryo lethality.

The successful generation of normal male and female offspring from flutamide treated PCOS-mice demonstrated that blockage of androgen receptor pathway prevents placenta dysfunction and supports proper fetal growth. This observation, coupled with previous findings indicating complete reversal of PCOS-like symptoms in both androgen receptor KO mice and flutamide-treated PCOS-mice, underscores the central role of androgen receptor pathways in the pathogenesis of PCOS, including the pregnancy complications.^[69, 70]^ Consistent with previous findings indicating that flutamide restores the endometrium receptivity,^[17]^ our study reveals comparable placental and liver transcriptional profiles between control mice and those treated with flutamide in the PCOS model, suggesting the prevention of hyperandrogenism-induced adverse events. However, it is noteworthy that flutamide treatment during pregnancy showed small adverse effects on male offspring, such as shorter AGD, in line with previous reports.^[71]^ In the context of women with PCOS, flutamide use is normally combined with other first-line treatment options to treat hirsutism related symptoms only.^[72]^ Thus flutamide should not be used during human pregnancy.^[47]^

## 4. Conclusion

All together, we revealed mid-gestation embryo lethality and deleterious effects on placental development in the androgenized mouse PCOS model. Notably, comparable adverse effects were observed in human trophoblast organoids, suggesting the presence of conserved effects induced by hyperandrogenism. All identified detrimental effects were effectively prevented by blocking androgen receptors, highlighting the crucial role of the androgen-mediated pathways in the pathogenesis of PCOS. These findings hold promise for future research aimed at discovering safe androgen receptor blockers and related strategies to mitigate the adverse impact of hyperandrogenism on pregnancy outcomes and offspring health.

## Acknowledgments

We thank Strategic Research Program in Diabetes at the Karolinska Institutet for the use of TSE Systems and EchoMRI in the Metabolic Phenotyping Centre; and the histological Core Facility-Histocore, Biomedicum at the Karolinska Institutet. This work is supported by Swedish Medical Research Council: project no. 2022-00550 (ESV), 2020-00253 (QD); Knut and Alice Wallenberg Foundation: 2019.0211 (QD); Distinguished Investigator Grant – Endocrinology and Metabolism, Novo Nordisk Foundation: NNF22OC0072904, and project grant NNF18OC0033992 and NNF19OC0056647 (ESV); Diabetes Foundation: DIA2021-633 (ESV); Karolinska Institutet KID funding: 2023-0005 and 2020-00990 (ESV); Karolinska Instiutet faculty funded position (QD); Regional Agreement on Medical Training and Clinical Research between the Stockholm County Council and the Karolinska Institutet: 20190079 (ESV).

## 5. Method

### Ethical approval

All animal experiments were approved by Swedish board of Agriculture (Jordbruksverket, ethical approval number: DNR20485-2020). Animal care and procedures were performed in accordance with guidelines specified by European Council Directive and controlled by Comparative Medicine Biomedicum (KM-B), Karolinska Institutet, Stockholm, Sweden. Human trophoblast stem cells were obtained from collected first trimester placenta due to elective pregnancy termination. The use of tissues and experimental procedures were approved by the Medical University of Vienna ethics boards (no. 084/2009).

### Animals and treatment

3-week-old C57Bl/6J female mice were purchased from Janvier Labs (Le Genest-Saint-Isle, France). Mice were housed five per cage in IVC-GM 500 cages with maintained temperature (22°C) and humidity (55–65%) at a 12/12 h light/dark cycle and fed with in house chow diet. DHT pellets were prepared according to previously published protocol. ^[26]^ Briefly, Dow Corning Silastic tubing 0.04 mm inner diameter × 0.085 mm outer diameter (Fisher Scientific, Hampton, NH) was filled with DHT (around 5.24 mg for 10mm DHT pellet or around 2.68 mg for 5mm DHT pellet, 5α-androstan-17β-ol-3-one; A8380, Sigma-Aldrich), and 2 mm medical adhesive silicone (Factor II, Lakeside, AZ) was used to seal the tube on both sides. For control pellets, 14 mm or 9 mm empty pellets were sealed directly. Pellets were incubated in saline for 24 hours at 37°C for equilibration before insertion.

At 4 weeks of age, mice were randomly divided into 3 groups: control, DHT (i.e. PCOS) and DHT with flutamide (DHT + FLU) i.e. PCOS and flutamide. In the PCOS group, a DHT pellet was implanted, and the control group received an empty pellet under light isoflurane anesthesia. Co-treatment with flutamide was done by implantation of a 90-day continuous-release pellet containing 25mg flutamide (Innovative Research of America, Sarasota, FL, USA) at the same time as the DHT pellet.

### Superovulation and mating

Four weeks after pellet implantation, mice from all groups were superovulated with 5IU pregnant mare’s serum gonadotropin (PMSG; Folligon, MSD Animal Health Care) followed by 5IU human chorionic gonadotropin (hCG; Pregnyl 5000IE, Merck Sharp & Dohme AB, Stockholm, Sweden) 48 hours later. Females were then mated with wild type, unexposed males overnight right after hCG injection, and the morning after mating was counted as embryonic day (E) 0.5. At embryonic day E10.5 and E13.5 pregnant females were dissected, and pregnancy was defined as >2g increase in body weight from E0.5.

### Reproductive phenotypic assessments

A separate batch of F_0_ mice was kept until 10-13 weeks of age, i.e. 7 weeks after pellet implantation, when their reproductive phenotypes were assessed. Estrus cyclicity was assessed by vaginal smear and cytology for 12 consecutive days as previously described.^[6]^ The anogenital distance (AGD) was assessed in F_0_ mice at the time of dissection. For F_1_ offspring, AGD was measured at around 16 weeks of age, then again at around 28 weeks of age. The estrus cyclicity was assessed twice when they reached around 4 months and 6 months of age with 12 days of assessment each time.

### Metabolic and behavioral phenotype assessments

The body weight of F_0_ mice was monitored throughout the experiment (from pellet implantation to 10 weeks after the treatment started). For F_1_ offspring, the body weight development was followed from the weaning of mice (3 weeks) if possible until the end of the study (dissection).

#### EchoMRI

Magnetic resonance imaging scans (EchoMRITM, Houston, TX, USA) was used to assess the body composition when they reached 4 and 6 months of age.

#### Basal metabolism

To measure the basal metabolism, mice were single housed in metabolic cages (TSE PhenoMaster, TSE Systems) for 3 consecutive days when they reached 4 months of age. Food intake, gas exchange and spontaneous locomotor activity were recorded by the cages every 3 minutes. The 24-hour reading started after the first adaptation day in the cage and used for analysis. RER was interpreted as the ratio between the volume of produced CO_2_ and the volume of consumed O_2_. EE was calculated by indirect gas calorimetry and adjusted for total body mass.

#### Oral glucose tolerance test (OGTT)

Mice were fasted for 6 hours prior to the OGTT; at the time of testing, tail blood was collected for measuring blood glucose with One tough ultra-2 glucometer at time 0, 15, 30, 60 and 90 mins after 20% glucose challenge. Blood was also collected before and 15 mins after glucose introduction to assess insulin levels with Ultra-Sensitive Mouse Insulin ELISA Kit 90080 (Crystal Chem Inc., Downers Grove, IL, USA).

#### Behavioral assessments

Elevated plus maze (EPM) and open field (OF) were used for testing behavior as previously described.^[19]^ All behavior experiments were conducted during the light phase, and mice were acclimatized in the testing room for at least 20 min prior to the test. In both tests, mice were tracked automatically by an infrared digital camera using the EthoVision XT Software (Noldus, Wageningen, The Netherlands).

### Tissue collection

At the end of phenotypic assessments, mice were subjected to tissue collection. Two hours fasting was performed before dissection, and tissue weight was recorded after collection. Tissues were snap frozen directly after collection in liquid nitrogen and stored in −80°C. The serum hormone measurement was performed with LC-MS/MS method as previously described.^[73]^

### Histological analysis of ovary and placenta

Ovaries and E10.5 and E13.5 placenta were dissected and fixed in 4% PFA overnight, dehydrated and embedded in paraffin. The ovary was sectioned at 8µm thickness and the placenta at 5µm and subjected for haematoxylin & eosin staining or stored at room temperature for later immunofluorescence (IF). One or two representative images were taken per section with a light microscope at ×10 magnification (Zeiss Axioplan).

### Immunofluorescence (IF)

#### Placenta

For IF staining, placenta sections were deparaffinized and rehydrated, then antigen retrieval was performed with citrate buffer (C9999, Sigma-Aldrich) for 6 mins in microwave (800W). After cooling down, sections were permeabilized with 0.1% Tween 20 then blocked with 5% donkey serum in PBS for 1 hour at room temperature. Then the sections were incubated overnight at 4°C with primary antibody (MCT1-Sigma ab1286-I, MCT4-Sigma ab3314P). The next day, sections were washed and incubated with secondary antibody (donkey anti-chicken IgG 647, Invitrogen; donkey anti-rabbit IgG 594, Invitrogen). Sections were then washed and incubated with DAPI then mounted with anti-fade mounting medium with DAPI (H-1800, Vectashield).

#### Human trophoblast organoid

Human trophoblast organoids were collected and fixed in 4% PFA overnight, they were then proceeded for dehydration with 30% sucrose (57501, Merck) for another 24 hours. Organoids were then washed and embedded in OCT for cryosectioning on Epredia NX70 cryostat. Sections are at 5µm thickness and stored in −80°C until use. At the time of IF staining, sections were let to sit at room temperature for 10 mins before start, then they were washed twice with PBS followed by blocking with PBST (5% BSA and 0.3% Triton-100 in PBS) for 2 hours at RT. Primary antibody (anti-ENDOU, HPA012388, Sigma-Aldrich; anti-EpCAM, VU1D9, Cell Signaling Technology) incubation was done overnight at 4°C followed by washing and 1 hour of secondary antibody (donkey anti-mouse IgG 488, Invitrogen; donkey anti-rabbit IgG 594, Invitrogen) incubation at RT. Sections were then washed and incubated with DAPI then mounted with anti-fade mounting medium with DAPI (H-1800, Vectashield). Confocal images were taken with Zeiss LSM800 confocal microscopy at 10X, 20X or 40X depending on the sample type.

### Embryo dissection and isolation of primordial germ cells

The placenta and embryos were dissected at E10.5 and E13.5 in ice-cold 1x PBS using a dissection microscope (VWR). At each time point, after dissection, the fetal crown–rump length and weight were measured, and embryos and placenta were imaged at 1.5x (E10.5) and 1x (E13.5) magnification. Embryos were subjected to PGC isolation and/or staining of PGCs using immunohistochemistry (IHC). For IHC: At E10.5, two whole embryos per dam were fixed and incubated with Anti-STELLA antibody to investigate germ cell migration by whole mount staining (see below) and images were acquired using a confocal laser-scanning microscope. For E13.5, the gonads from 1 male and 1 female embryo per dam was fixed in 4% PFA overnight and subjected to IHC for PGC counting. For PGC isolation, the central body containing the hindgut of the remaining E10.5 embryos were dissected, dissociated and FACS sorted to separate somatic and germ cells using SSEA-1 antibody and the integrin beta 1 (CD61) for WGB and RNA sequencing (see below). ^[74]^ The gonads from each E13.5 embryos were dissected and sex determinate before they were pooled together based on sex and dam for FACS sorting and WGB and RNA sequencing (see below) ^[74]^. The gonads from each E13.5 embryos were dissected and sex determinate before they were pooled together based on sex and dam for FACS sorting and WGB and RNA sequencing (see below).

*Whole mount staining* of the E10.5 embryos were performed according to previously established protocols with modification. ^[75]^ Briefly, three E10.5 embryos from each group were fixed in 4% PFA on rotation at 4°C for 2 hours with rotation, followed by rising with TPBS (PBS + 0.3% Triton-X) at 4 ° C for three times at 1 hour interval. Then the samples were incubated with rabbit anti-STELLA (1:1000, Abcam) in TPBS for at 4°C overnight with rotation. This was followed by rinsing the samples overnight in TPBS at 4°C with rotation. On the next day, the samples were incubated with Donkey anti-rabbit IgG conjugated with Alexa488 in TPBS (1:2000, Life Technologies) overnight at 4°C with rotation. Samples were then washed in TPBS overnight at 4°C with rotation and stored in dark at 4°C until the clearing process. Tissue clearing was performed by washing the samples at room temperature sequentially in 50% Methanol/PBS for 5 mins with rotation, 3 times in 100% Methanol for 20 mins, and lastly in BABB (benzyl alcohol/benzyl benzoate; Sigma Aldrich) for 5 mins. Samples were further cleared in BABB for around 1 hour at RT with rotation until it became completely transparent. Cleared samples were immediately proceeded for confocal microscopy (Zeiss LSM880).

#### Microscope

Whole mount cleared embryos were transferred to glass bottom dishes (MatTec) containing BABB for imaging with Zeiss LSM880 confocal microscope.

#### Image processing

For whole mount-stained embryos, the stacked images were first converted to imaris file using ImarisFileConverter and analyzed using Imaris x64 software (version 9.5.1, Bitplane). The surface tool and mask function were used to crop out region of PGC in the whole embryo. Then spot function was applied to the cropped region to identify PGCs. Number of PGCs for each embryo was used for further analysis.

#### Dissociation and FACS sorting

The specimens for FACS were dissociated in dissociation buffer containing 1mg/mL Collagenase (Sigma C0130) + 0.1mg/mL DNaseI (Roche 11284932001) in PBS at 37°C for 6 mins with slow shaking. After that the specimens were further dissociated by pipetting in washing buffer containing 0.1% BSA (Sigma A7030) + 0.1mg/mL DnaseI in PBS on ice. Then the cell suspension was centrifuged at 220g for 5 mins and the cell pellets rinsed twice with washing buffer. The resulting pellets were resuspended in 200 µL of chilled washing buffer and incubated with PE-conjugated anti-CD61 antibody (1:200, BioLegend 104307) and eFluor660-conjugated anti-SSEA1 antibody (1:20, eBioscience 50-8813-42) for 15-30 mins on ice. After that, the cells were rinsed twice with washing buffer. Prior to cell sorting, the cell suspension was further dissociated by passing through a 35um cell strainer (Corning 352235) and stained with cap Fixable viability Dye fluor450 (eBiscience 65-0863-14) or Dapi. FACS sorting was performed with a 100 mm nozzle using a flow cytometer (SONY SH800S) following to the manufacturer’s instructions. Embryonic germ cells (SSEA+CD61+) and somatic cells (SSEA-CD61-) were sorted directly into Smartseq3 lysis buffer for downstream processing for RNA sequencing and whole genome bisulphite sequencing and stored at −80°C until use.

### Human trophoblast organoid generation

The human trophoblast stem cell lines were generously given by our collaborator Sandra Haider. They werecultured and developed into organoids according to previous publications.^[48]^ Briefly, trophoblast stem cells were cultured in Fibronectin (Sigma-Aldrich FC010) coated plate, and the culture medium was Advanced DMEM/F12 (Gibco 11540446) supplemented with 1x B-27^TM^ Supplement (50X) minus vitamin A (Gibco 12587010), 1x Insulin Transferrin Selenium Ethanolamine (ITS-X) (Gibco 10524233), 1x L-glutamine(Thermo Fisher 25030024), 1µM A83-01 (Tocris 2939), 3µM CHIR99021 (Tocris 4423), 50ng/ml hEGF (Gibco PHG0311), 5µM Y-27632 (Sigma Aldrich Y0503), 0.1 mg/ml Gentamicin (Fisher Scientific 15-710-064) and 0.01M HEPES (Gibco 15710-049). Once the trophoblast stem cells reach 80-90% confluency, they were seeded into Matrigel to form trophoblast organoids. Detailed procedure of organoid formation was performed according to previous publications. Briefly, 7-8000 total trophoblast stem cells were mixed with 20 µl of Matrigel and this Matrigel mix drop was placed onto a pre-warmed 48-well plate. The plate was put in the incubator for 15 minutes before the addition of 250µl of trophoblast medium. The trophoblast medium composition was as follows: advanced DMEM/F12, 1x B27 without vitamin A, 1x ITS-X, 1x L-glutamine, 0.1mg/ml Gentamicin, 0.01M HEPES, 1uM A83-01, 3µM CHIR99021, 100ng/ml hEGF, 5µM Y-27632. For experimental purposes to mimic a hyperandrogenic environment, the following 2 mediums were prepared based on the trophoblast medium: DHT medium with the addition of 1nM DHT and FLU medium with the addition of 1nM DHT and 2uM Flutamide. For the prolactin experiments, the addition of 100ng/ml prolactin was added to the trophoblast medium with or without the supplementation of DHT. The medium was changed every 2-3 days until the majority of the organoids in control medium reach 200 mm in diameter.

#### Differentiation of human trophoblast organoid

To assess the ability of trophoblast stem cells to differentiate into extravillous trophoblasts, we stimulated the differentiation with the following differentiation medium. Diff1: advanced DMEM/F12, 1x B27 without vitamin A, 1x ITS-X, 1x L-glutamine, 0.05mg/ml Gentamicin, 0.01M HEPES, 2uM A83-01, 50ng/ml hEGF. Diff2: advanced DMEM/F12, 1x B27 without vitamin A, 1x ITS-X, 1x L-glutamine, 0.05mg/ml Gentamicin, 0.01M HEPES, 50ng/ml hEGF. When the majority of the organoids reach the size of 200 mm, Diff1 was used to stimulate differentiation for 5 days, after 5 days, the organoids were wash with washing medium (composition: advanced DMEM/F12, 1x B27 without vitamin A, 1x ITS-X, 1x L-glutamine, 0.05mg/ml Gentamicin, 0.01M HEPES) then supplemented with Diff2 medium for another 5 days before they were harvested for RNA extraction. For experimental purpose for mimicking the hyperandrgenism and study the effect of flutamide, the following differentiation medium were prepared: DHT Diff1 and DHT Diff2 mediums with the supplementation of 1nM DHT, and Flu Diff1 and Flu Diff2 mediums with the supplementation of 1nM DHT and 2uM flutamide.

#### hCG measures

At the end of each experiment, hCG was measured in each plate with commercial pregnancy test strips (Gravidtetstest GI29100, Medistore).

#### Human trophoblast organoids size measurement

At the end of experiments, pictures were taken for the organoids using microscope at 4X magnification with a scale. Images were processed with CellProfiler (version 4.0.7) for organoid identification and size measurement.

### Smart-Seq3 bulk RNA sequencing library preparation

cDNA libraries for placenta and PGCs were generated using the Smart-seq3 protocol.^[76]^ cDNA libraries for placenta and PGCs were generated using the Smart-seq3 protocol. ^[76]^ At E10.5 were 10-50 PGCs per dam sorted into lysis buffer, visual sex determination was performed at E13.5 and 5-50 PGCs/sex was sorted into lysis buffer for each dam for Smart-seq 3 (**Supplementary Table 8**). Genotyping was performed to determine the sex of embryos from E10.5 and E13.5, and RNA from placenta of both E10.5 and E13.5 were extracted using Trizol method. The placenta RNA was pooled for sequencing according to sex and dam, except for DHT group at E10.5 stage where only 1 dam was pregnant, and the placenta were sequenced individually. RNA was reverse transcribed using Maxima H minus reverse transcriptase (Thermo Fisher). Resulting cDNA was amplified with KAPA HiFi HotStart ReadyMix (KAPA Biosystems) by 18 cycles of PCR, then libraries purified with 22% PEG Clean-up beads. Quality check was performed using an Agilent 2100 BioAnalyser (Agilent Technology) to assess quality and quantity of the cDNA library. 100pg cDNA each sample was tagmented using a Tn5 transposase and amplified for 8 cycles using Nextera Index Primers (Ilumina Novaseq 6000).

### Smart-Seq3 bulk RNA sequencing mapping and quantification

Raw demultiplexed fastq files were merged and processed using zUMIs^[77]^ (v2.9.7c) with STAR^[78]^ (2.7.10a) to generate expression profiles. Reads were mapped against GRCm38 and were quantified with gene annotations from GENCODE GRCm38.p6.

### Prime-Seq bulk RNA sequencing library preparation

cDNA libraries for embryo liver were generated using the Prime-Seq protocol. ^[79]^ E13.5 liver RNA was isolated using a combination of TRIReagent (Sigma) and ReliaPrep RNA Miniprep Systems (Promega). Then liver RNA were pooled for sequencing according to sex and dam. The extracted RNA was normalized to a concentration to 10 ng/uL, and 40ng in total was used from each sample to provide the library. RT was performed using Maxima H Minus Reverse Transcriptase (ThermoFisher Scientific), followed by preamplification using KAPA HiFi 2X ReadyMix (Roche). Thereafter, the obtained cDNA was normalized to a concentration of 6 ng/uL. Subsequent steps were performed using the NEBNext Ultra II FS DNA Library Prep Kit (New England Biosciences), as described in the protocol for prime-seq. Agilent Bioanalyzer 2100 High Sensitivity DNA Analysis Kits (Agilent) were used to QC both the cDNA and final library quality and fragment size.

### Prime-Seq bulk RNA sequencing mapping and quantification

The adapters and polyG from Raw non-demultiplexed fastq files were trimmed using cutadapt (v4.1) and fastp (0.23.2). The clean reads were processed using zUMIs^[77]^ (v2.9.7c) with STAR^[78]^ (2.7.10a) to generate expression profiles. To extract and identify the UMI-containing reads in zUMIs, the base_definition: BC(1–12) and UMI (13– 28) were specified for file 1 and cDNA (15–150) for file 2 in the YAML file. UMIs were collapsed using a Hamming distance of 1. Reads were mapped against GRCm38 and were quantified with gene annotations from Gencode GRCm38.p6.

### Gene expression analysis

From the RNA quantification matrices, we filtered out the low-expressed genes. Genes were removed if they were expressed in the number of samples less than the average group size, i.e., total number of samples divided by group numbers. Principal component analysis and Spearman correlation were calculated based on log2 normalized variance stabilized transformed counts.

For differentially expressed analysis, the RNA quantification matrices were analyzed using DESeq2^[80]^ with the treatment condition as the variable of interest. Statistically significant genes (log2 foldchange >1, adjusted P < 0.05) were identified and then used for making plots and for downstream analysis.

For Cell type deconvolution analysis, the single nuclei^[51]^, or single cell^[28]^ RNA count matrix from E10.5 placenta with the annotated cell type information and our RNA-seq count matrix was processed using the CIBERSORTx^[81]^ to impute the placental or trophoblast cell type fractions.

### Whole genome bisulfite sequencing library preparation

At E10.5 50-200 PGCs per dam were sorted into lysis buffer, sex determination was performed at E13.5 and 25-500 PGCs/sex were sorted into lysis buffer for each dam for WGBS (***Supplementary Table 6***). Genotyping was performed to determine the sex of embryos from E10.5, and DNA from placenta of both E10.5 and E13.5 was extracted using QIAamp Fast DNA kit (Qiagen). Then placenta DNAs were pooled for sequencing according to sex and dam, except for DHT group at E10.5 stage where only 1 dam was pregnant, and the placenta were sequenced individually. 1µg of genomic DNA each sample was spiked with 6.54 ng unmethylated lambda DNA then sonicated into a mean size of 350bp fragments with ME220 Focused-Ultrasonicator (Covaris). The ends of sonicated DNA fragments were repaired before adaptor ligation with NEBNext Ultra II DNA Library Prep kit for Ilumina (New England BioLabs) according to manufacturer’s instruction. The adaptor ligated DNA was cleaned up with 0.8x AMPure xp (Beckman Coulter) beads to remove any adaptor dimer or small fragments. The ligated DNA was bisulfite converted with the EZ DNA Methylation-Gold kit (Zymo Research) then PCR amplified for 10 cycles with KAPA HiFi Hotstart Uracil+ kit (Roche). 2% agarose gel was used to perform size selection of the amplified products and QIAquick Gel Extraction kit (Qiagen) to extract product of desired size.

### Whole genome bisulfite sequencing data processing

For WGBS data, the illumina and library specific adapters were trimmed using bbduk (v38.98, BBMap - Bushnell B. - sourceforge.net/projects/bbmap/) with parameters ‘ktrim=r k=23 mink=11 hdist=1 tpe tbo qtrim=rl trimq=10 minlen=2’. The reads after trimming were mapped to mouse genome GRCm38, deduplicated and methylated Cs were called using bismark.^[82]^ Coverage was calculated as ‘read length x number of uniquely mapped reads / 2652783500’ using a customed script. Differentially methylated sites and regions were analyzed using MethylSig package in R.^[83]^ Further annotation was made using AnnotationHub package In R.

### Analyzing embryonic and placental development phenotype-relevant transcriptome changes

For testing the co-association for the androgen excess induced embryonic and placental development phenotypes relevant gene expression changes, we first retrieved the Mammalian Phenotype Ontology for all the differentially expressed genes (in PCOS-mice placenta at E10.5 compared to Control-mice). The enrichment analysis was performed on up- or down-regulated gene sets using Fisher’s exact test and the p-values were adjusted using the Benjamini-Hochberg method for correction for multiple hypotheses testing. The co-association was inferred from whether the down-regulated genes by androgen excess in the given Mammalian Phenotype Ontology significantly overlap. This was tested using Fisher’s exact test.

### Statistical analysis

Two-way repeated measure ANOVA was used for body weight development and analysis was performed in SPSS. For all F_0_ group comparisons were one-way ANOVA with Turkey post hoc test or Kruskal-wallis test with Dunn’s test used. All F_0_ statistics were done using GraphPad Prism (version 8.4.3) for Windows. For all embryo and F_1_ group comparisons, analysis of covariance (ANCOVA) was used whenever possible to adjust for litter. ANCOVA statistics was performed in R (version 4.2.1). Data are presented as mean values ± s.e.m. Differences were considered statistically significant when P < 0.05.

## Data availability

The data that support the findings of this study are available within the paper in Supplementary Information files and source data are provided in Mendeley Data, V1, doi: 10.17632/hb9n3jxr4c.1. The raw sequence data reported in this paper have been deposited under the Bioproject accession no. PRJNA1001956.

## Author contribution

H.L. designed the study, performed the mouse data collection, performed molecular analyses, analysed the data, prepared the figures, prepared the sequencing library and wrote the manuscript. H.J. analysed the RNA and whole genome bisulfite sequencing data, prepared the figures, and was involved in manuscript preparation. C.L. was involved in the study design, preparation of the sequencing library, interpretation of the results, and preparation of the manuscript. E.D. was involved in the human organoid planning and revising the manuscript. A.Z was responsible for prime seq library preparation. G.E. was involved in mouse experiment. HP.P. was involved in the embryo collection. S.R. performed offspring molecular analysis. Y.P. was involved in the RNA sequencing library preparation. T.M. was involved in preparing human trophoblast stem cells. C.O. performed the serum sex steroid analyses with GC-MS. E.L. performed the mouse embryo image data analyses. S.H. was involved in preparing human trophoblast stem cells and revising the manuscript. A.B., was involved in the study design, interpretation of the results, and preparation of the manuscript; E.S-V. and Q.D. designed the study, analysed the data, prepared the figures, and wrote the manuscript. All authors read and approved the final version of the manuscript.

## Competing interests

No authors have any conflict of interest to declare.

---

**Extended Data Fig. 1.**
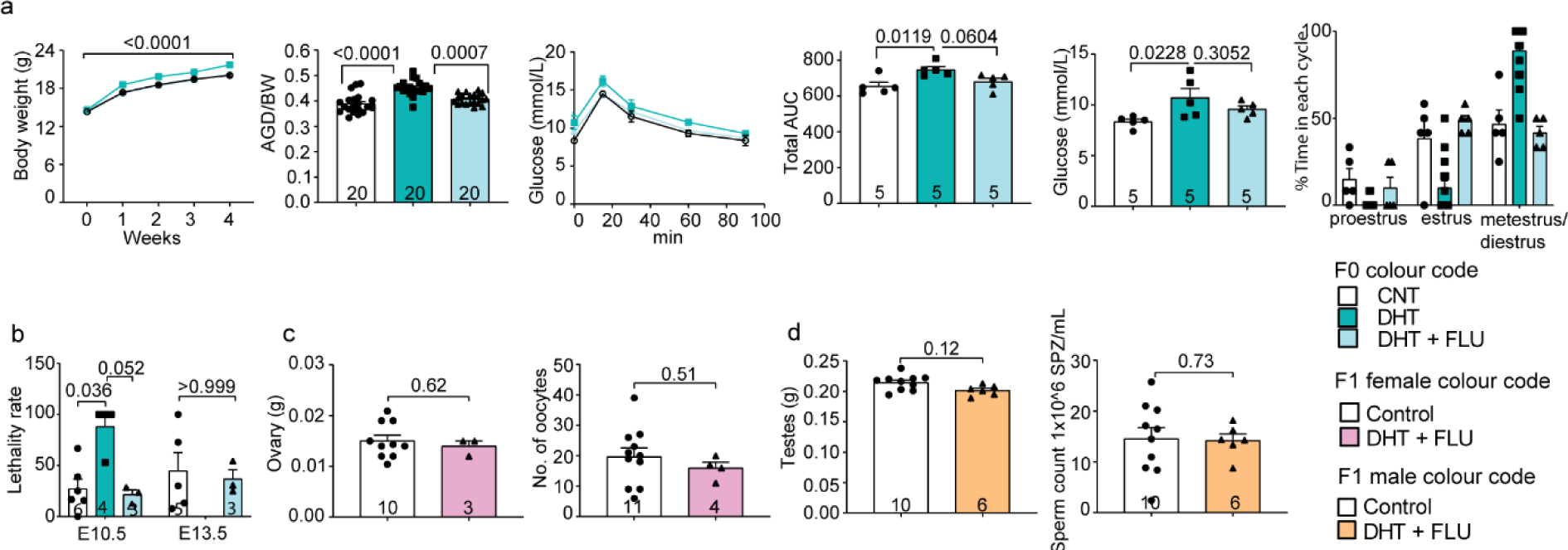
Phenotypes of peripubertal F0 dams with reduced dose and F1 offspring. **a,** 5mm DHT-implanted PCOS-mice phenotyping, including: body weight development (CNT = 25, DHT = 40, DHT + FLU = 25), AGD normalized by body weight, glucose tolerance and basal glucose level, and estrus cyclicity after 4 weeks of implantation. **b,** Total number of dead/absorbed embryos/dam at the time of dissection. **c,** F1 female offspring ovary weight and number of oocytes per donor after dissection. **d,** F1 male offspring testis weight and number of motile sperms after dissection. Numbers of mice are stated in the bars of each group. Body weight development: Repeated measures ANOVA. Statistics: F0 statistics are done with One-way ANOVA with Turkey post hoc analysis; F1 statistics performed by ANCOVA to control for litter.

**Extended Data Fig. 2.**
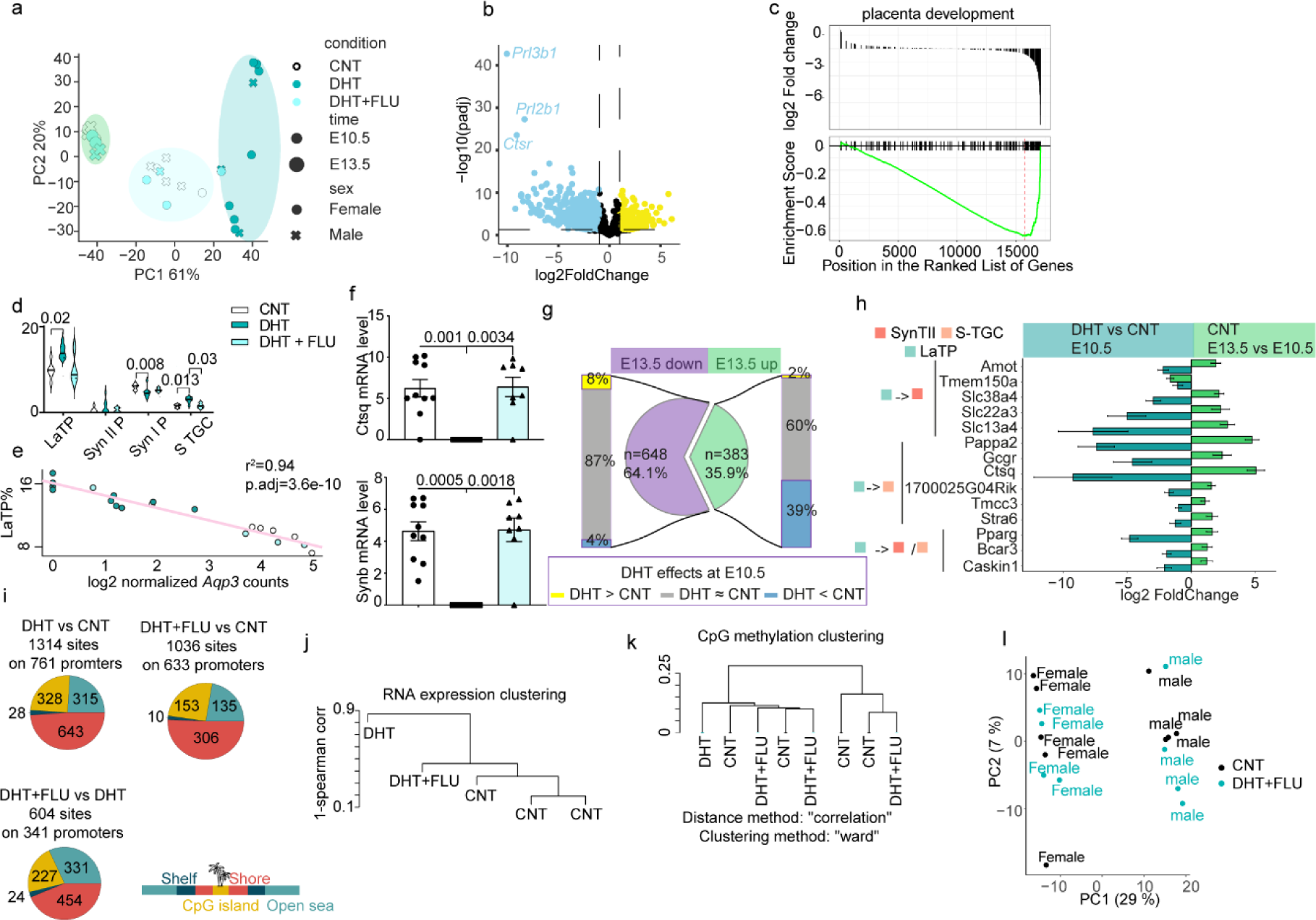
Flutamide prevent the major RNA expression and DNA methylation changes in placentae and offspring livers induced by excess androgen exposure. **a,** The first two principal components of gene expression matrices from the placenta samples. **b**, Vocalbo plot of the gene expression from placentas at E10.5 comparing DHT with Control group. **c,** Gene set enrichment analysis reveals negative enrichment in placenta development in the placenta of DHT-implanted PCOS-mice compared to control mice. **d,** Cell type proportion deconvoluted aided by the single cell expression atlas for trophoblast at E10.5**. e,** Linear regression of selected genes with labyrinth trophoblast precursors. P values are adjusted using the Holm method. **f,** Log normalized transcript counts of *Ctsq* in E10.5 placenta. **g,** The overlap between DEGs comparing DHT treated PCOS-mice and control mice at E10.5 or at E13.5. **h,** DEGs varied along the differentiation trajectory. The bar plot shows the fold change of genes at E10.5 or E13.5 by DHT treatment. The genes were the DEGs comparing DHT treated PCOS-mice and control mice at E10.5 or DEGs comparing E13.5 to E10.5. The genes were annotated with the differentiation trajectory if they were varied in the respective pseudo time trajectory is from. LaTP: labyrinth trophoblast progenitor, S-TGC: sinusoid trophoblast giant cells. **i,** Pie plots representing differentially methylated promoters’ distribution on CpG annotations. **j,** Clustering with the distance metrics 1-spearman correlation from the gene expression in E10.5 PGC. **k,** Clustering of E10.5 PGC samples based on CpG methylation. **l,** The first two principal components of gene expression matrices from the liver samples collected at 6 months in F1 generation. In **d**, **e** and **i** data are represented as mean ± s.d.

## Notes

### Competing Interest Statement

The authors have declared no competing interest.

